# TBX15 regulates a network of immune response genes in adipose tissue and alters fat mass and depot weight in heterozygous knockout mice

**DOI:** 10.1101/2024.09.20.614167

**Authors:** L. Zolkiewski, M. Simon, J. Harrison, L. Vizor, E. Ireson, L. Moir, M. Yon, L. Beresford, A. Rodrigues, S. Hill, J. Hawkins, L. Bentley, R. D. Cox, R. Dumbell

## Abstract

Adipose tissue distribution in the body is an indicator of metabolic disease risk, independent of body mass index (BMI), and is indirectly measured by waist-hip-ratio (WHR). T-Box transcription factor-15 (TBX15) has been implicated in regulation of adipose distribution in multiple human and mouse studies, and the *TBX15-WARS2* genome-wide association study locus has been associated with BMI-adjusted-WHR signals in multiple investigations. As a potential mediator of this signal, we investigated the role of *Tbx15* using heterozygous and homozygous mouse knockout models to determine if loss of this gene alters adipose physiology, and to identify the transcriptional network regulated by *Tbx15* in adipose tissue and preadipocyte cells. In a metabolic phenotyping experiment we provided either low fat diet (LFD) or high fat diet (HFD) to male and female heterozygous *Tbx15^+/-^* and wildtype *Tbx15^+/+^*mice from weaning and maintained for 24 weeks. Only *Tbx15^+/-^*mice maintained on LFD weighed less than wildtype LFD controls, and female LFD *Tbx15^+/-^* mice had lower fat mass overall. We found that in LFD *Tbx15^+/-^* mice, multiple visceral fat depots weighed less than wildtype controls, and this was maintained when corrected for body mass for both gonadal and mesenteric visceral adipose depots. When comparing adipocyte size in multiple adipose depots, some reduction in number of larger adipocytes was detected in the perirenal adipose tissue of female HFD *Tbx15^+/-^* vs *Tbx15^+/+^* mice, mesenteric adipose tissue from female LFD *Tbx15^+/-^* vs *Tbx15^+/+^* mice and male HFD *Tbx15^+/-^* vs *Tbx15^+/+^*mice. RNA-sequencing of subcutaneous (inguinal) adipose tissues from 12-week old male and female knockout *Tbx15^-/-^*, *Tbx15^+/-^* and *Tbx15^+/+^* mice raised on a standard chow diet identified 897 upregulated genes and 2328 downregulated genes in female *Tbx15^-/-^* mice compared to *Tbx15^+/+^*mice. We then combined this dataset with TBX15 ChIP-sequencing data from mouse preadipocyte 3T3-L1 cells overexpressing *TBX15* to identify a *credible set* of genes directly regulated by TBX15. These 52 genes were enriched for B- and T-cell receptor signalling, JAK-STAT signalling and haematopoietic cell lineage pathways; suggesting a direct regulatory role for TBX15 in these pathways in adipose tissue. Together, these data highlight a role for TBX15 in regulation of differential adipose tissue expansion, particularly under low caloric conditions. Further, we identify a potentially important role for TBX15 in the well described adipocyte-immune cell crosstalk associated with obesity and type 2 diabetes mellitus.

## Introduction

Obesity is associated with increased risk of cardiovascular disease, type 2 diabetes mellitus (T2DM), metabolic syndrome and reduced life expectancy. Waist-hip ratio (WHR) is an anthropomorphic measure indicating abdominal adiposity that coupled with inadequate expansion of subcutaneous depots, is a stronger predictor of metabolic disease than BMI alone ^1–6^. White adipose tissues (WAT) are heterogeneous populations of mature adipocytes, mesenchymal stem cells, endothelial and smooth muscle cells, while also containing immune cells such as macrophages, innate lymphoid cells and T-cells. WAT expansion occurs through hyperplasia and/or hypertrophy of existing adipocytes, coupled with an acute pro-inflammatory response and extracellular matrix remodelling ^7–9^. WAT expansion in obesity is characterised by expression of pro-inflammatory cytokines and recruitment of “M1” macrophages resulting in low-grade chronic inflammation associated with insulin resistance, obesity and ultimately T2DM ^10^.

Genome-wide association studies (GWAS) have identified thousands of single nucleotide polymorphisms (SNPs) associated with WHR independent of overall BMI (WHRadjBMI) ^11–14^. Signals in the *TBX15-WARS2* locus are well replicated in these studies, however, there appears to be a complex regulatory network at play with evidence of sex differences in several signals within the locus ^11–13^. We have previously investigated the *Wars2* gene in this locus in multiple mouse and *in vitro* models and found that a hypomorphic allele shows pleiotropic effects that include reduced adiposity and adipose tissue dysfunction, sex and diet specific effects on fat distribution and browning of white adipose tissue, with some of these effects mediated by elevated FGF21 and GDF15 ^15,16^.

T-box transcription factor-15 (TBX15) is a transcriptional regulator whose gene expression in adipose tissue was inversely correlated with WHRadjBMI, insulin, HOMA-β and triglyceride levels and positively associated with fat-free mass and Matsuda Index, among other cardiometabolic traits ^17^. TBX15 is predominantly expressed in skeletal and muscle cells, with more recent *in* work highlighting an increased fat mass in heterozygous knockout *Tbx15* mice fed a high-fat diet (HFD) ^18,19^. Further, *Tbx15* is expressed at high levels in subcutaneous adipose tissue and to a lesser extent in visceral adipose tissue ^20^. *In vitro, Tbx15* both activates and represses adipogenesis genes in mouse 3T3-L1 and primary sub-vascular fraction (SVF) cells and has been implicated in browning of white adipocytes ^19–22^.

Thus, we aimed to investigate the role of TBX15, as one of the potential effector genes within the *TBX15-WARS2* WHRadjBMI locus, in adipose tissue and body composition in mice, and to define a transcriptional network regulated by TBX15 in mouse adipose tissues and human and mouse (pre-)adipocyte cell lines. Here, we show that *Tbx15* heterozygous knockout mice exhibit lower fat mass vs wildtype controls when a fed a low-fat diet. Further, using RNA- and ChIP-sequencing we highlight a potential transcriptional regulatory role for TBX15 in immune response signalling in WAT.

## Materials and Methods

### Animals

Mice were maintained following UK Home Office legislation and local ethical guidelines issued by the Medical Research Council (Responsibility in the Use of Animals for Medical Research, July 1993). Procedures were approved by the UKRI MRC Harwell Animal Welfare and Ethical Review Board (AWERB). Mice were maintained at the Mary Lyon centre (MLC) at MRC Harwell, with 12h light and dark cycle with a temperature of 21±2°C and 55±10% humidity. Frozen *Tbx15* heterozygous (*Tbx15^+/-^*) embryos were purchased from the Jackson laboratory (JAX:000551-B6EiC3 a/A-Tbx15de-H/J). Embryos were implanted into CD1 females to generate F_0_ litters. Male and female *Tbx15^+/-^* mice were subsequently backcrossed to C57BL/6NTac mice.

Male and female *Tbx15^+/-^* mice at backcross 2 were used to generate intercross litters for an initial viability cohort (n= 35 mice total; one male *Tbx15* ^-/-^ mouse was culled due to small size) and RNA-sequencing cohort (n=27 mice total). Male *Tbx15^+/-^*mice were backcrossed to female C57BL/6NT mice to generate phenotyping cohorts at backcross 5, wild-type (*Tbx15^+/+^*) colony mates were used as controls (n=153 mice total; one male *Tbx15^+/-^*mouse was culled due to malocclusion). . Incomplete data from culled mice were not included in statistcal evaluation or presentation of data. Genotyping was performed on ear-clips at 2 weeks of age by the MRCH genotyping facility. Mice were randomised to single sex, mixed genotype cages at 3 weeks of age and all phenotyping was performed blinded to genotypes.

### Experiment 1 (metabolic phenotyping cohort)

For metabolic phenotyping cohorts, mice were weaned and randomised, with *ad libitum* access to food, either low-fat diet (10 kcal% fat; D12450J, Research Diets Inc., New Brunswick, NJ, USA) or high-fat diet (60 kcal% fat; D12492, Research Diets Inc.), and water (25 ppm chlorine). Power calculations and biologically significant difference estimation were calculated on data collected from the initial viability cohort (n=35 mice) using GPower3 software. Based on these calculations, an n=20 animals per treatment group was considered sufficient to detect biological differences. Therefore, a total of 153 mice were weaned across three metabolic phenotyping cohorts (Female HFD *Tbx15^+/+^* n=18, female HFD *Tbx15^+/-^* n=22, female LFD *Tbx15^+/+^* n=18, female LFD *Tbx15^+/-^* n=21, male HFD *Tbx15^+/+^* n=21, male HFD *Tbx15^+/-^* n=17, male LFD *Tbx15^+/+^*n=21, male LFD *Tbx15^+/-^* n=15; one male LFD *Tbx15^+/-^* mouse was culled due to malocclusion). Mice were metabolically phenotyped up to 25 weeks of age.

### Experimental 2 (tissue RNA-sequencing cohort)

Mice for RNA-sequencing cohorts were weaned and randomised with *ad libitum* access to food (rat and mouse no. 3 breeding diet; RM3) and water (25 ppm chlorine). A total of 27 mice were weaned into one cohort. Mice were aged to 12 weeks before tissue collection for RNA-sequencing.

### Metabolic phenotyping

Total body mass was measured every two weeks from 4 weeks of age on a scale calibrated to 0.01g. Body composition (total fat mass (g) and total lean mass (g)) was measured every two weeks using an Echo-MRI (Echo-MRI-100, Echo-MRI, Texas, USA).

Mice were fasted for 16-18 hours (at 12-13 weeks of age for intraperitoneal glucose tolerance tests; IPGTT) and 4-5 hours (at 17 weeks of age for intraperitoneal insulin sensitivity test; IPIST). Subsequently, total body mass (g) was measured on a scale calibrated to 0.01g and EMLA^TM^ cream (5% lidocaine/prilocaine, cat. PL39699/0088, Aspen Pharma Trading Ltd, Dublin, Ireland) was applied to tails 15 minutes prior to starting the test. A small incision was made in the tail vein and baseline blood glucose levels (0 minutes) were measured using an AlphaTrak2 glucose monitor with AlphaTrak2 test strips (cat. ART2406-003 Zoetis, Parsippany-Troy Hills, NJ, USA).

For IPGTT, 20% (w/v) glucose solution was administered intraperitoneally (final dose 2 g/kg bodyweight). Blood glucose levels were further measured at 30, 60 and 120 minutes after glucose administration. For IST, hypurin porcine insulin neutral solution (FU40177, Wockhardt Ltd., Mumbai, India) was diluted in NaCl solution and administered intraperitoneally at a final dose of 1.25 IU/kg (HFD males), 1.0 IU/kg (LFD males and HFD females) or 0.5 IU/kg (LFD females). Blood glucose levels were further measured at 15, 30, 45, 60 and 90 minutes after insulin administration.

For terminal procedures, 24–25-week-old mice were fasted for 4-5 hours. Mice were euthanised by inhalation overdose of isoflurane. Once fully anaesthetised, blood was collected into lithium-heparin microvette tubes (078028, KABE Labortechnik GmbH, Nümbrecht, Germany) from the retro-orbital sinus and centrifuged at 5000 *x g* for 10 minutes at 4°C. Plasma was transferred to a clean microcentrifuge tube and stored at -80°C. Mice were dissected for tissues including visceral fat depots: gonadal (gWAT), mesenteric (mWAT), perirenal (pWAT) and the subcutaneous inguinal (iWAT) white adipose depot, interscapular brown adipose tissue (BAT) and liver. All tissues were weighed on a scale calibrated to 0.0001 g and were fixed in 10 % formalin (cat. 38008010C, Leica, Wetzlar, Germany) for ≥ 24 h at a minimum 20x volume. Terminal blood plasma samples collected from metabolic phenotyping cohorts were measured by the MLC clinical pathology laboratory on a DXC 700 AU clinical chemistry analyser (Beckman Coulter, High Wycombe, UK). Samples were run in batches grouped by sex and diet.

### Adipocyte size comparisons in multiple adipose tissues

Fixed tissues from a subset of mice (n=4 per sex and diet; n=32 total mice) were sent to Histologix Ltd. (Nottingham, UK) for processing and staining. Tissues were embedded in paraffin wax, sectioned (WAT = 8 μM; BAT and liver = 5 μM), and stained with haematoxylin and eosin (H&E) following standard protocols. Per mouse, 1x section per gWAT, mWAT, pWAT and iWAT were scanned at 20X magnification on a NanoZoomer RS with NDP Scan v3.2 software (Hamamatsu, Hamamatsu, Japan). Up to 10 regions of interest were exported at 10x magnification using the NDP.view2 v2.9.29 Software (Hamamatsu). Cell area for WAT samples was analysed using the Adiposoft plugin in ImageJ, with the following calibration settings following initial optimisation: microns per pixel 0.920, minimum diameter 15, and maximum diameter 150 ^23,24^. Adiposoft analysis was run in automated batch mode and data was exported to Microsoft Excel for average cell area and cell size distribution analysis.

### RNA-Sequencing

A further cohort of male and female *Tbx15^+/+^* and *Tbx15^-/-^* mice raised on standard chow diet (SDS Rat and Mouse No. 3 Breeding diet, RM3; n=27 mice total) were euthanised at 12 weeks old by cervical dislocation. iWAT and BAT tissues were collected and frozen in LN_2_ and stored at --70°C. RNA was isolated from frozen adipose tissues using the RNeasy Lipid Tissue Mini Kit (74804, Qiagen, Hilden, Germany) following manufacturer’s instructions. RNA quality was determined using the Agilent RNA 6000 Nano Kit (5067-1511, Agilent Technologies, Santa Clara, CA, USA) and chips were read using the 2100 software and a Bioanalyzer (Agilent Technologies). Samples with a RNA Integrity Number (RIN) above 7.0 were considered high enough quality for subsequent sequence analysis. RNA yield, 260/280 and 260/230 ratios were determined using a Nanodrop ND-8000 (ThermoFischer Scientific, Waltham, MA, USA) and 1000 ng RNA was plated per well of a 96-well plate for RNA-Seq workflows. Library preparation, RNA sample processing and generation of initial sequencing data were performed by the Oxford Genomics Centre.

### ChIP-Sequencing

TBX15 cDNA constructs (602 amino acids) were designed and purchased from Origene Technologies. The TBX15 sequence was subcloned into a C-terminal tagged HA vector (636590, TaKaRa)_by PCR amplification (Q5 polymerase, M0491, NEB; forward primer *AATGTCGACGCCGCCGCGATCGCCATGA;* reverse primer *AATGGTACCAACCATGTGCACGGACATCT*), restriction enzyme digestion (SalI-HF, RS3138, NEB and KpnI-HF RS3142, NEB; 37°C for 60 minutes, followed by 65°C for 20 minutes) and ligation (T4 DNA ligase M0202, NEB), following manufacturers’ standard protocols. Murine 3T3-L1 cells were purchased from American Type Culture Collection (cat. ATCC-CL-173, lot 61194648, ATCC, Manassas, VA, USA) and were maintained in DMEM-GlutaMAX^TM^ supplemented with 10% (v/v) newborn calf serum (NBCS; 260101-074) and 1% (v/v) penicillin-streptomycin. Cells were transfected with 46 µg HA-tagged TBX15 cDNA constructs per T175 flask using the Lipofectamine 3000^TM^ kit (L3000-015, Invitrogen), following manufacturer’s protocols. 24 hours later, cell culture samples were fixed and pelleted according to protocols from Active Motif Inc., snap frozen and shipped on dry ice to Active Motif Inc.. Active Motif Inc. prepared chromatin, performed chromatin immunoprecipitation (ChIP) reactions, generated and sequenced libraries on an Illumina NextSeq 500, and performed basic data analysis for ChIP-sequencing.

### NGS analysis

RNA-sequencing was aligned using STAR aligner ^25^ using GRCm38 (mm10) reference genome. Annotations were performed using htSeq ^26^ with Ensembl v79 ^27^. Differential genes between mutant and wildtype mice were determined using EdgeR ^28^. Functional analysis was performed using gProfiler. Basic ChIP-sequencing analysis was performed by Actif Motif Inc. ChIP-sequencing data was aligned to the GRCm38 (mm10) reference genome using the Burrows-Wheeler aligner algorithm ^29^. Peak locations were determined using the model-based analysis of ChIP-sequencing (MACS) algorithm v2.1.0, with a cut-off p<1×10^−7^ ^30^. Signal maps and peak locations were used as input data to Active Motif’s proprietary analysis program, which creates Excel tables containing detailed information on sample comparison, peak metrics, peak locations, and gene annotations.

### Phenotypic Data Statistical analysis

All *in vivo* data were analysed using GraphPad Prism version 9.0. Graphs show mean ± standard deviation, all test details are described in the figure legends. For continuous data, statistical comparisons were carried out by repeated measures 2-way ANOVA, separated for diet and sex with Holm-Šídák *post-hoc* comparisons. Tolerance test data were normalised to the value at 0 minutes, and compared by 2-way ANOVA, separated for diet and sex with Holm-Šídák *post-hoc* comparisons, and area under the curve was calculated for each individual mouse and compared by unpaired t-test within diet and sex. For final tissue weights, comparisons were by 2-way ANOVA, separated by diet, with Holm-Šídák *post-hoc* comparisons within sex. Adipocyte areas were compared by multiple t-test per bin / depot / sex / diet. P<0.05 was considered statistically significant, unless stated otherwise.

## Results

### *Tbx15^+/-^* and *Tbx15^-/-^* breeding viability

Homozygous knockout *Tbx15* (*Tbx15^-/-^*) mice were markedly smaller than wild-type *(Tbx15^+/+^)* littermates and exhibit significantly reduced viability at weaning (p=0.0350 [χ_2_ test] n= 2, 22 and 10, homozygote, heterozygote and wildtype respectively; data not shown), consistent with previous observations ^31^. Therefore we did not continue to phenotype *Tbx15^-/-^* mice in any long-term experiments. Heterozygous knockout *Tbx15 (Tbx15^+/-^)* mice were born at expected Mendelian ratios and were indistinguishable from *Tbx15^+/+^* littermates at birth, however, in male mice there were significantly fewer *Tbx15^+/-^* mice born than *Tbx15^+/+^* colony mates (p=0.0295 [***x***_2_ test]; n= 32 and 53 respectively, data not shown).

### *Tbx15^+/-^* mice have lower body mass and body composition

Male and female *Tbx15^+/-^* and *Tbx15^+/+^* mice at 3 weeks of age were weaned onto a high fat (HFD; 60% fat), or low fat (LFD; 10% fat) diet and we assessed metabolic phenotypes up to 25 weeks of age. To assess whether *Tbx15* affects body composition, we measured body composition using EchoMRI every 2 weeks (Figure ***1***A-F). Male and female mice were compared separately for statistical analysis. For both male and females, *Tbx15^+/-^*mice weighed less than wildtypes when maintained on a LFD (p = 0.0226 and 0.0326, respectively), and no significant effects of genotype were detected on body mass for HFD mice (Figure ***1***A, D). In female LFD mice there was significantly lower fat mass in *Tbx15^+/-^* mice (p = 0.0329), and an interaction between this effect and time (p = 0.0011; Figure ***1***B). No significant effects of genotype on fat mass were detected in male mice, however an interaction between genotype and time was detected in LFD male mice (p = 0.0014). This was not accompanied by any significant post hoc time-point tests (Figure ***1***E). Similarly, significant interactions were detected between genotype and time for both LFD and HFD female mice lean mass (p = 0.0281 and 0.0162, respectively; Figure ***1***C) with no significant post hoc tests. No significant differences were detected for lean mass in any comparisons made in male mice (Figure ***1***F).

**Figure 1.**
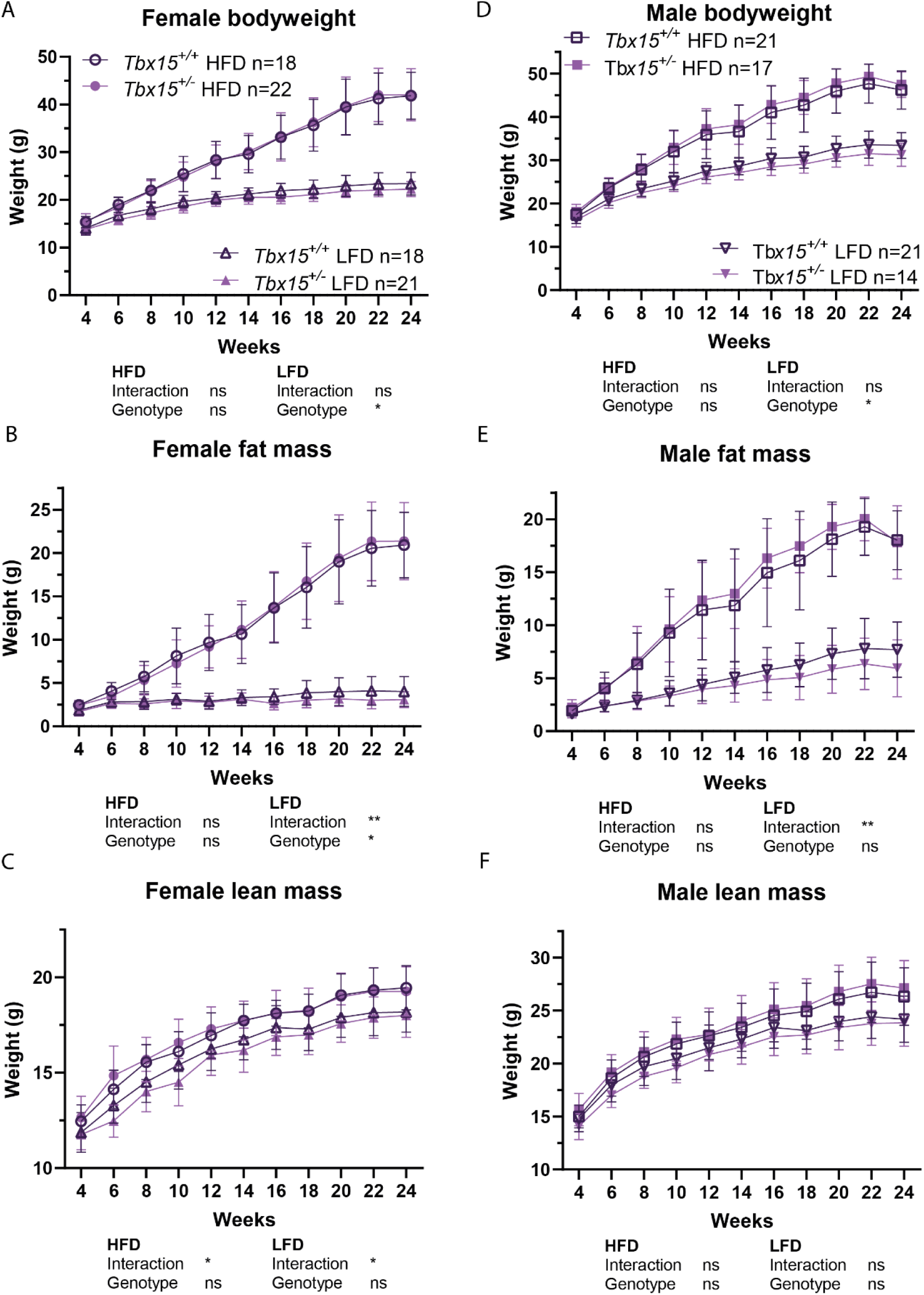
Low-fat diet *Tbx15^+/-^*mice have altered body composition at 4 to 24 weeks of age vs wildtype littermate controls. Female (A-C; n=18-22 per group), male (D-F; n=14-21 per group). A, D. Bodyweight. B, E. Fat mass. C, F. Lean mass. *Tbx15^+/+^*(dark purple), *Tbx15^+/-^* (purple), female high-fat diet (HFD; circle), female low-fat diet (LFD; upwards triangle), male HFD (square), male LFD (downwards triangle). Data are presented as mean ± standard deviation. Statistical significance was calculated by repeated measures ANOVA for overall genotype and interaction (time x genotype) effects, with Holm-Šídák multiple comparisons test within each diet: *p<0.05, **p<0.01. HFD, high-fat diet; LFD, low-fat diet.

### Female *Tbx15^+/-^* mice are less glucose tolerant when maintained on LFD

LFD female *Tbx15^+/-^* mice had a significantly decreased glucose tolerance profile compared to LFD *Tbx15^+/+^* (p = 0.0403 Figure 2A). However, no significant differences in glucose tolerance evaluated by area under the curve or fasted (t=0) glucose were detected in female mice fed a HFD or male mice fed a HFD or LFD (Figure 2). No significant differences were detected in blood glucose levels in response to insulin challenge in either HFD or LFD male or female mice (Figure 3).

**Figure 2.**
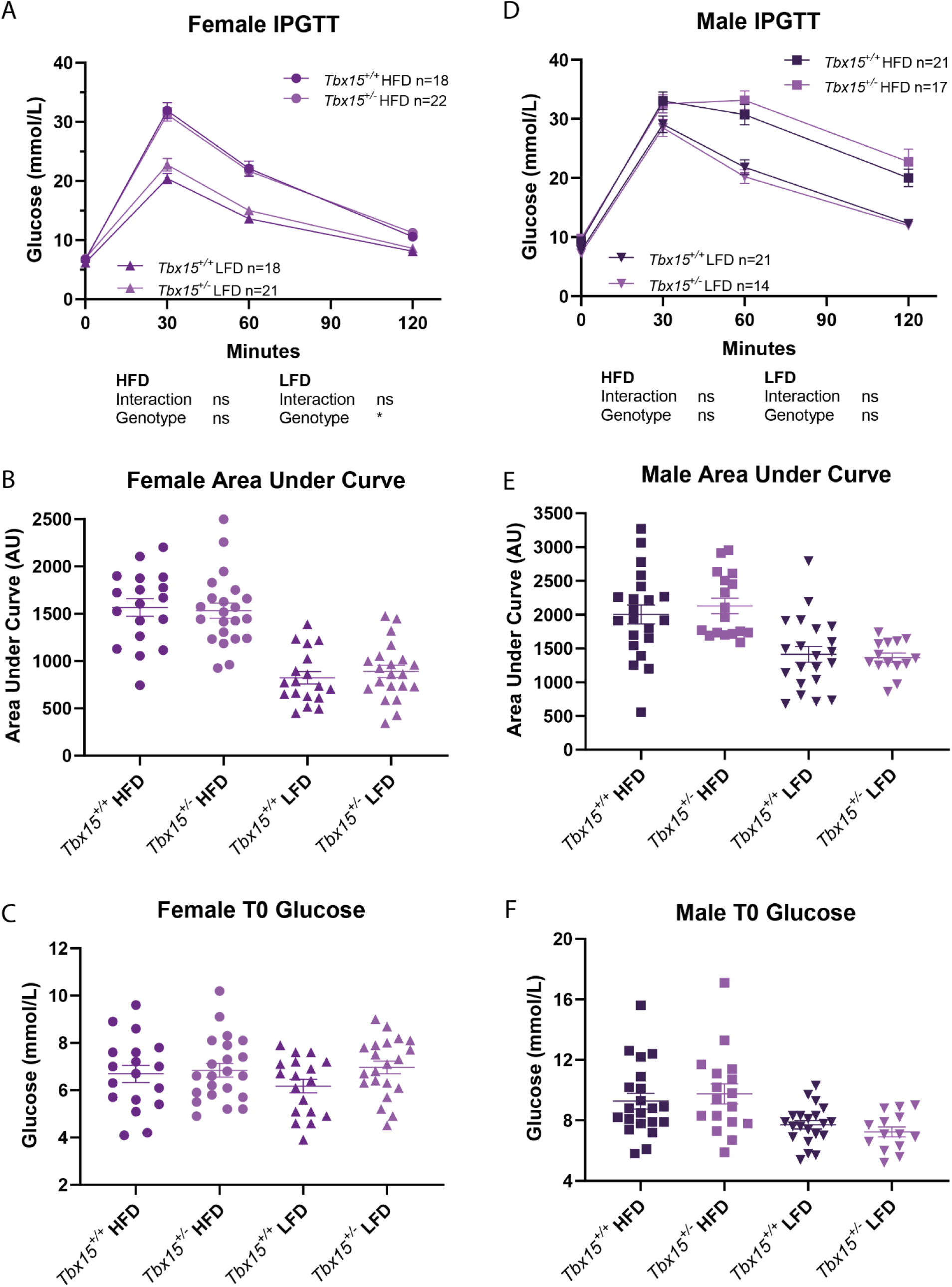
*Tbx15^+/-^*intraperitoneal glucose tolerance tests (IPGTT) at 13 weeks. Female (A-C; n=18-22 per group), male (D-F; n=14-21 per group). A, D. IPGTT. B, E. Area under curve. C, F. Fasted blood glucose (T0). *Tbx15^+/+^*(dark purple), *Tbx15^+/-^*(purple), female high-fat diet (HFD; circle), female low-fat diet (LFD; upwards triangle), male HFD (square), male LFD (downwards triangle). IPGTT data was analysed by two-way ANOVA with post-hoc Šídák’s multiple comparisons, interaction (time x genotype), genotype: *p<0.05. Area under curve and T0 glucose data was analysed by unpaired t-tests.

**Figure 3.**
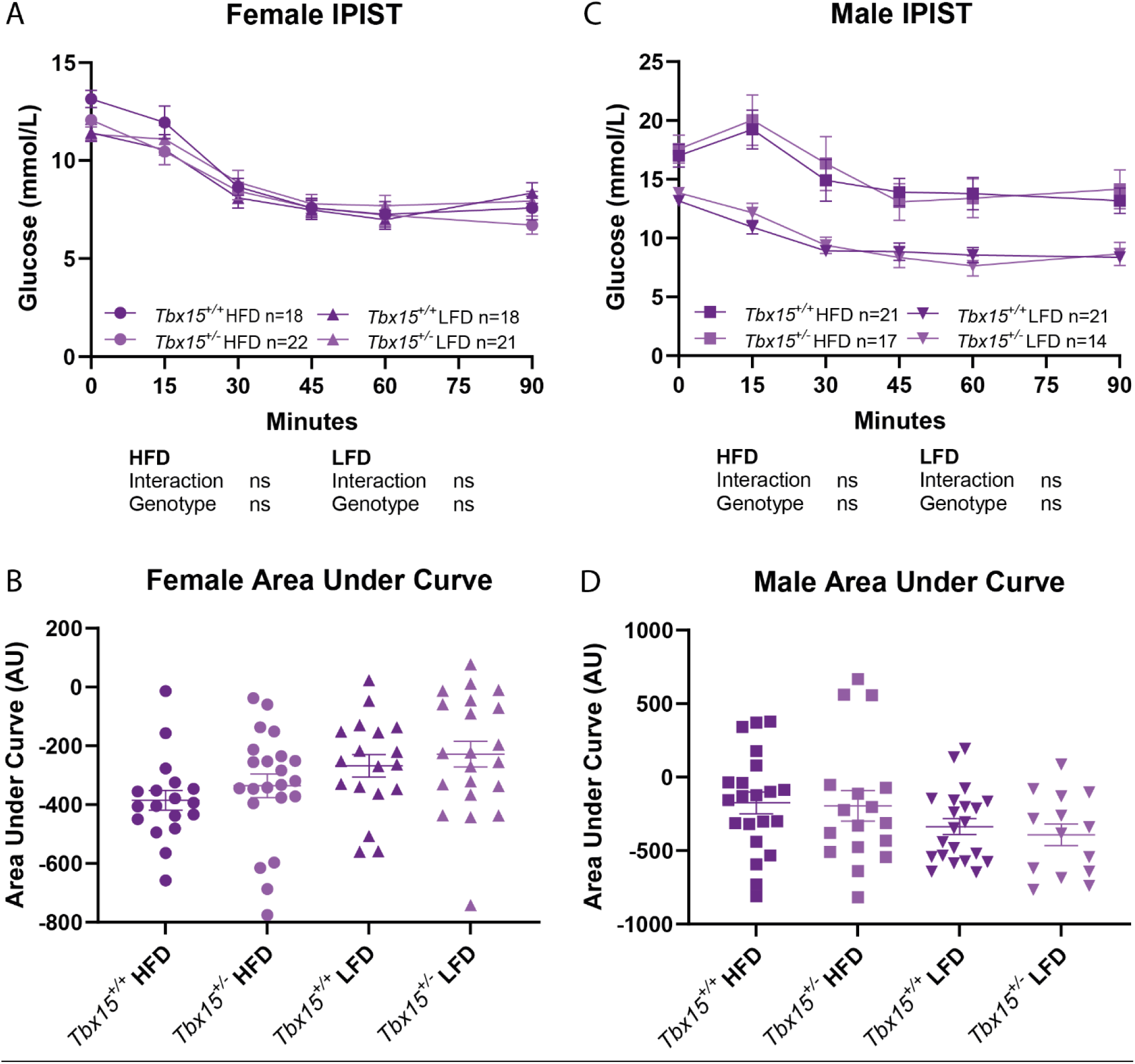
*Tbx15^+/-^* intraperitoneal insulin sensitivity test (IPIST) data at 17 weeks. Female (A-B), male (C-D). A, C. IPIST. B, D. Area under curve (AUC). *Tbx15^+/+^* (dark purple), *Tbx15^+/-^* (purple), female high-fat diet (HFD, circle), female low-fat diet (LFD, upwards triangle), male HFD (square), male LFD (downwards triangle). IPIST data analysed by two-way ANOVA with post-hoc Šídák’s multiple comparisons per sex and diet. Interaction (time x genotype). Area under curve data was analysed by unpaired t-tests per sex and diet.

### Terminal adipose tissue weights are lower in *Tbx15^+/-^* mice maintained on LFD

Next, we aimed to assess whether differences in adipose depot distribution contributed to the overall bodyweight and fat mass differences between *Tbx15^+/+^* and *Tbx15^+/-^* mice fed LFD or HFD.

Overall, at 24-25 weeks of age mice on HFD did not demonstrate any significant differences in bodyweight or individual fat depot or liver weights between for either male or female mice of either genotype. However, when adjusted for bodyweight (BW), overall pWAT/BW was lower for heterozygous mice (p = 0.0440), and in female mice had significantly increased gWAT /BW (adjusted p = 0.0154). We then compared gWAT : iWAT ratio as an indicator of visceral : subcutaneous fat distribution (as in WHR) and found no significant differences for either diet, genotype or sex combination (Table 1).

**Table 1.**
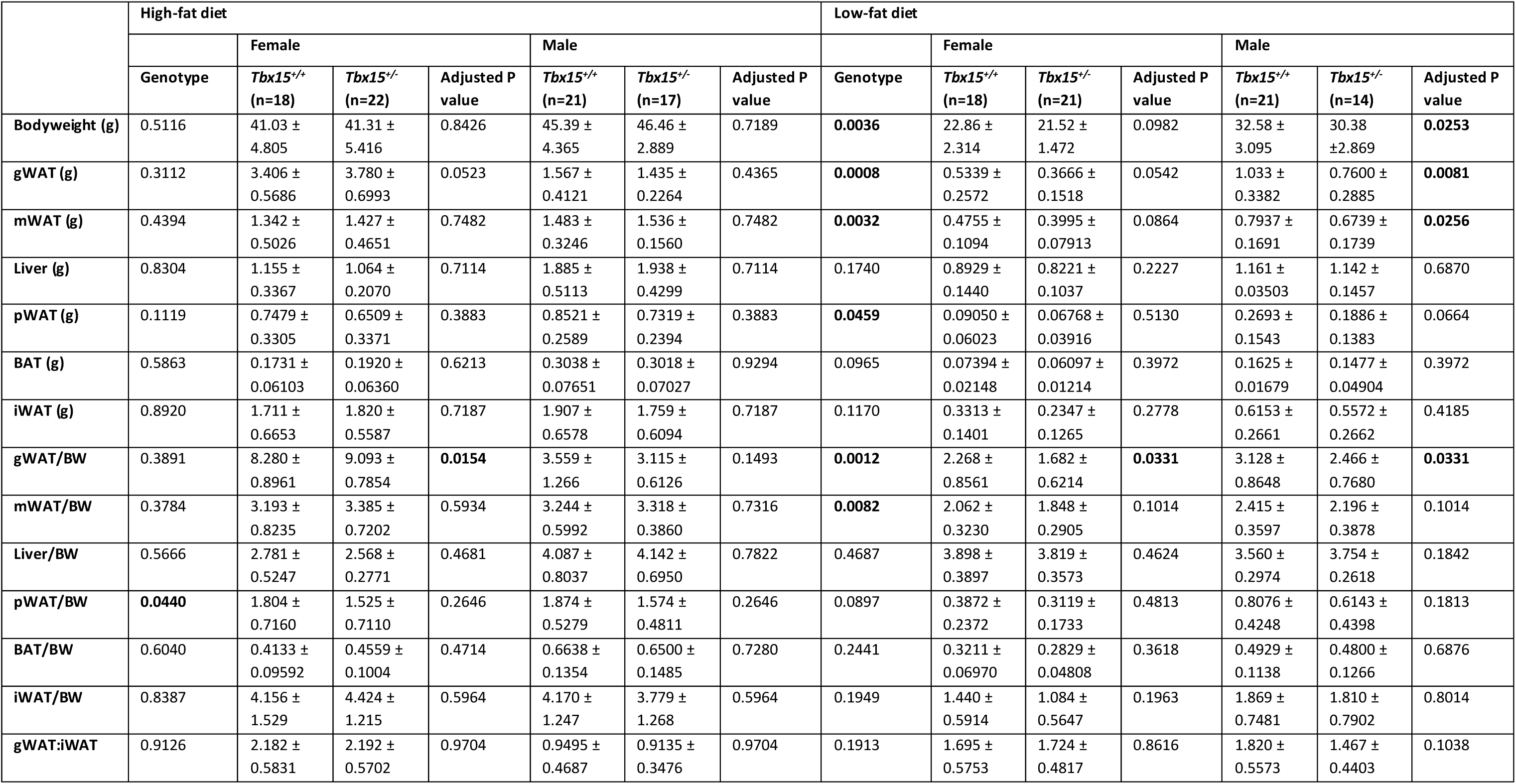
*Tbx15^+/-^* depot weights at 24 weeks of age. Data are presented as mean ± standard deviation (n=14-22 per group). Statistical significance was calculated for each diet group by two-way ANOVA for overall genotype and interaction (sex x genotype) effects, with Holm-Šídák multiple comparisons test within each sex (adjusted p value): significant values (p<0.05) are shown in bold. Data are shown as mean ± standard deviation. BAT, brown adipose tissue; BW, bodyweight; /BW, adjusted for bodyweight; gWAT, gonadal white adipose tissue, iWAT, inguinal white adipose tissue, mWAT, mesenteric white adipose tissue, pWAT, perirenal white adipose tissue, SD, standard deviation.

LFD mice, *Tbx15^+/-^* mice had significantly lower bodyweight (p = 0.0036), gWAT (p = 0.0008), mWAT (p = 0.0032) and pWAT (p = 0.0459) depots weights, with differences in gWAT (p = 0.0012) and mWAT (p = 0.0082) remaining statistically significant when adjusted for bodyweight (Table 1). When within-sex differences were compared, we found significantly lower bodyweight (adj. p = 0.0253), gWAT (adj. p = 0.0082) and mWAT (adj. p = 0.0256) in male *Tbx15^+/-^*compared to *Tbx15^+/+^* on LFD. When corrected to bodyweight, both female and male *Tbx15^+/-^* LFD mice had lower gWAT/BW (both adj p = 0.0331; Table 1).

### Altered adipocyte size distribution in fat tissues from *Tbx15^+/-^* mice

We then investigated whether there were differences in the adipocyte cell size within different WAT depots. For HFD female WAT, there were fewer larger adipocytes in *Tbx15^+/-^* mice in gWAT (3000-3999 µm; p = 0.039792) and pWAT (3000-3999 µm and 5000-9999 µm [in 1000 µm bins]; p = 0.040590, 0.043859, 0.039851, 0.012670 and 0.017420) Figure 4A, C); no significant differences were detected in adipocyte area distribution in mWAT or iWAT (Figure 4 B, D). Similarly, in HFD males there were fewer larger adipocytes in *Tbx15^+/-^* mice (7000-11999 µm [in 1000 µm bins] and 13000-13999 µm; p = 0.031032, 0.025291, 0.041103, 0.015381, 0.003100 and 0.010542) in mWAT ;; and more smaller adipocytes (2000-2999 µm; p = 0.010307) in iWAT (Figure 5B, D). No significant differences were detected in adipocyte area distribution in LFD female or male WAT (Figure 4 and Figure 5, respectively), with the exception of LFD *Tbx15^+/-^* female mice, who had fewer larger adipocytes in mWAT ( 5500-5999 µm; p = 0.030400; Figure 4F). This suggests an alternative mechanism, such as altered cell numbers rather than differences in adipocyte cell size, may be driving the altered fat mass and adipose depot distribution phenotypes observed in these mice.

**Figure 4.**
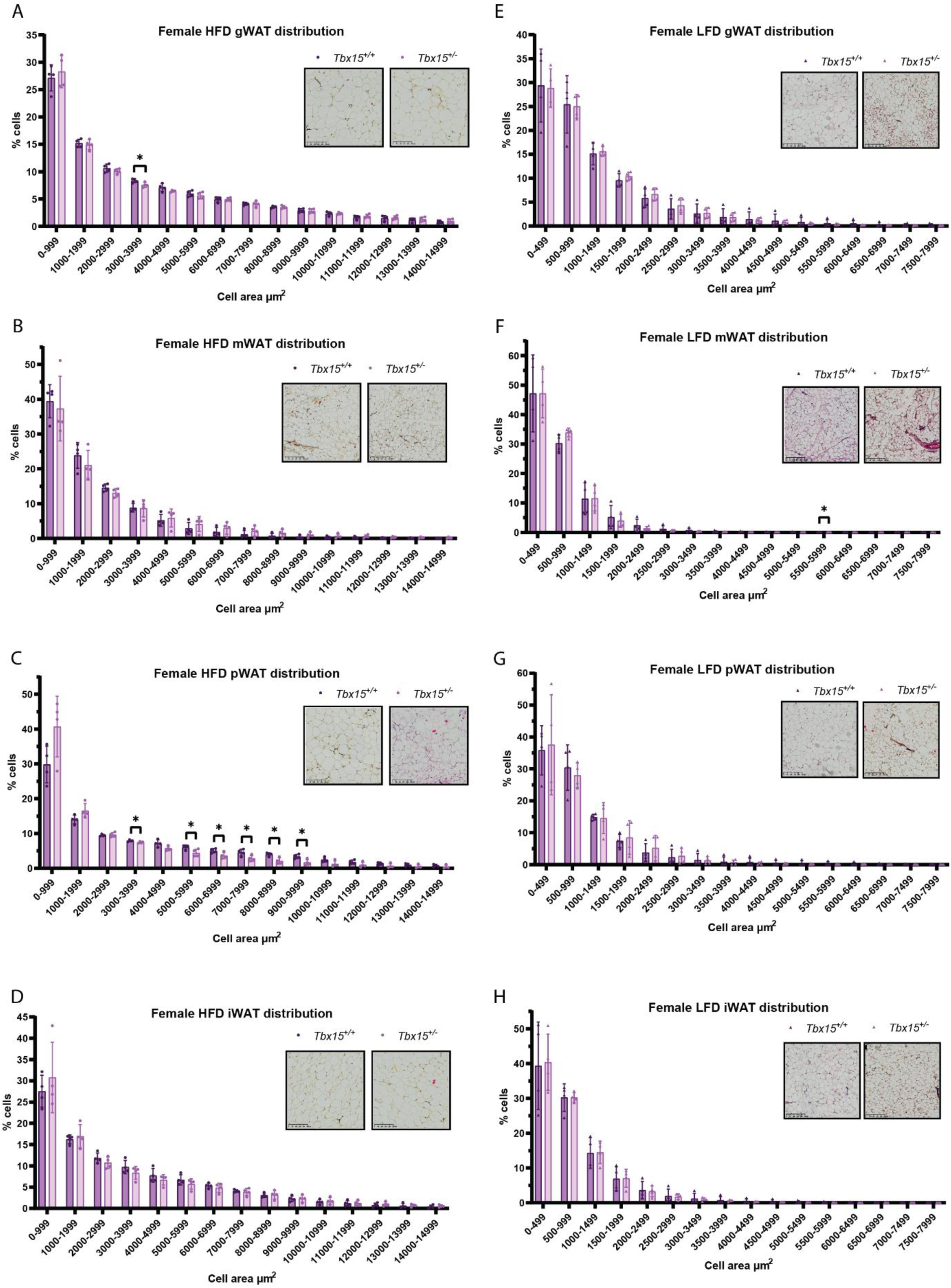
High-fat diet female *Tbx15^+/-^* mice have altered distribution of adipocyte cell areas at 24 weeks of age. **High** High-fat diet (A-D), low fat diet (E-H). A, F. gonadal WAT (gWAT). B, F. inguinal WAT (iWAT). C, G. mesenteric WAT (mWAT). D, H. perirenal WAT (pWAT). *Tbx15^+/+^* (dark purple), *Tbx15^+/-^* (purple), female high-fat diet (HFD; circle), female low-fat diet (LFD; upwards triangle). Data are presented as mean ± standard deviation (n=4 per group). Statistical significance was calculated by multiple unpaired t-tests within each sex, diet and depot: *p<0.05.

**Figure 5.**
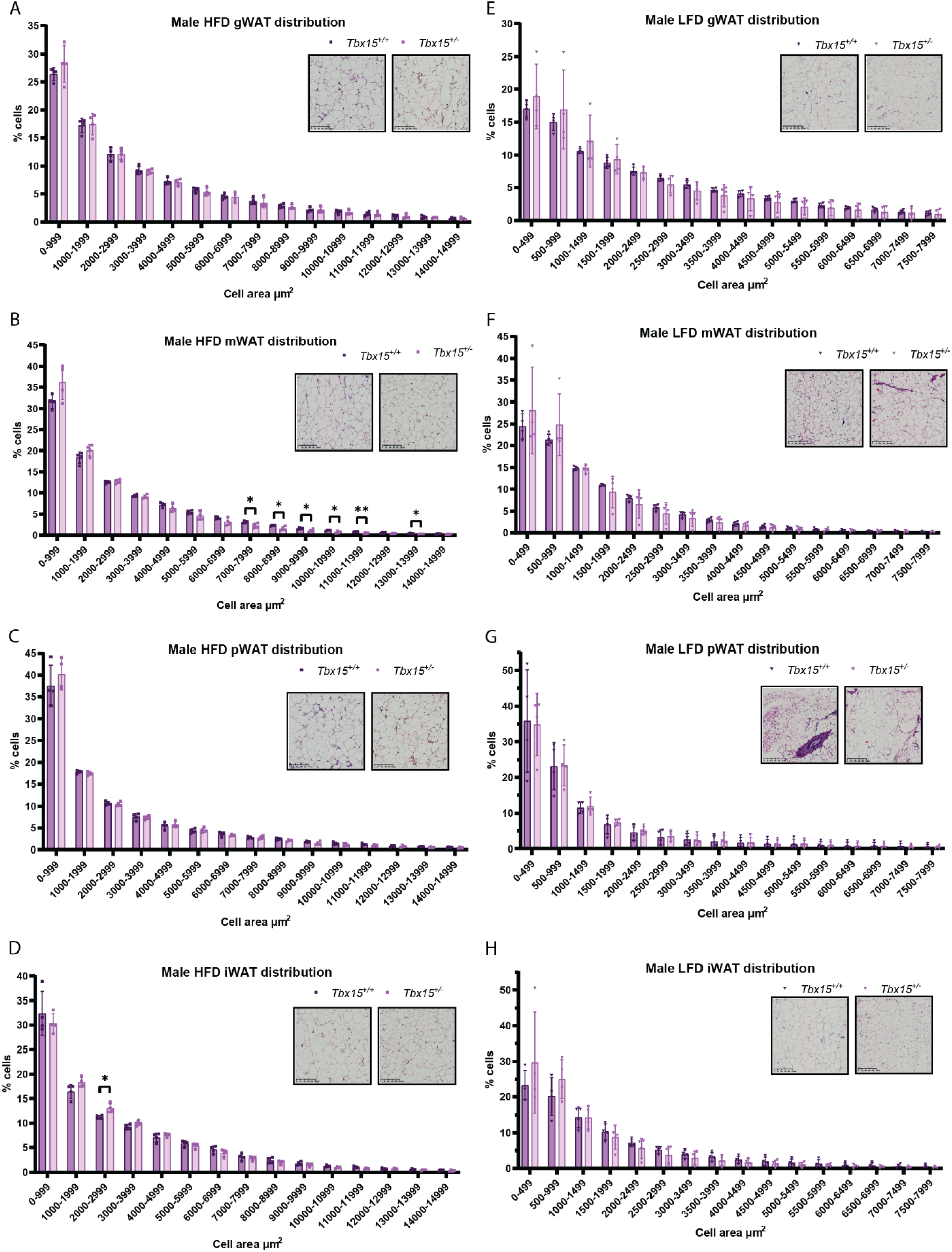
High-fat diet male *Tbx15^+/-^*mice have altered distribution of adipocyte cell areas at 24 weeks of age. High-fat diet (A-D), low fat diet (E-H A, F. gonadal WAT (gWAT). B, F. inguinal WAT (iWAT). C, G. mesenteric WAT (mWAT). D, H. perirenal WAT (pWAT). *Tbx15^+/+^*(dark purple), *Tbx15^+/-^*(purple), male HFD (square), male LFD (downwards triangle). Data are presented as mean ± standard deviation (n=4 per group). Statistical significance was calculated by multiple unpaired t-tests within each diet: *p<0.05, **p<0.01.

### Plasma analytes did not differ in *Tbx15^+/-^* mice

When comparing the 27 plasma analytes in both HFD and LFD male and female mice that might explain differential body compositions we found no effect of heterozygous knockout of *Tbx15* (Table 2).

**Table 2.**
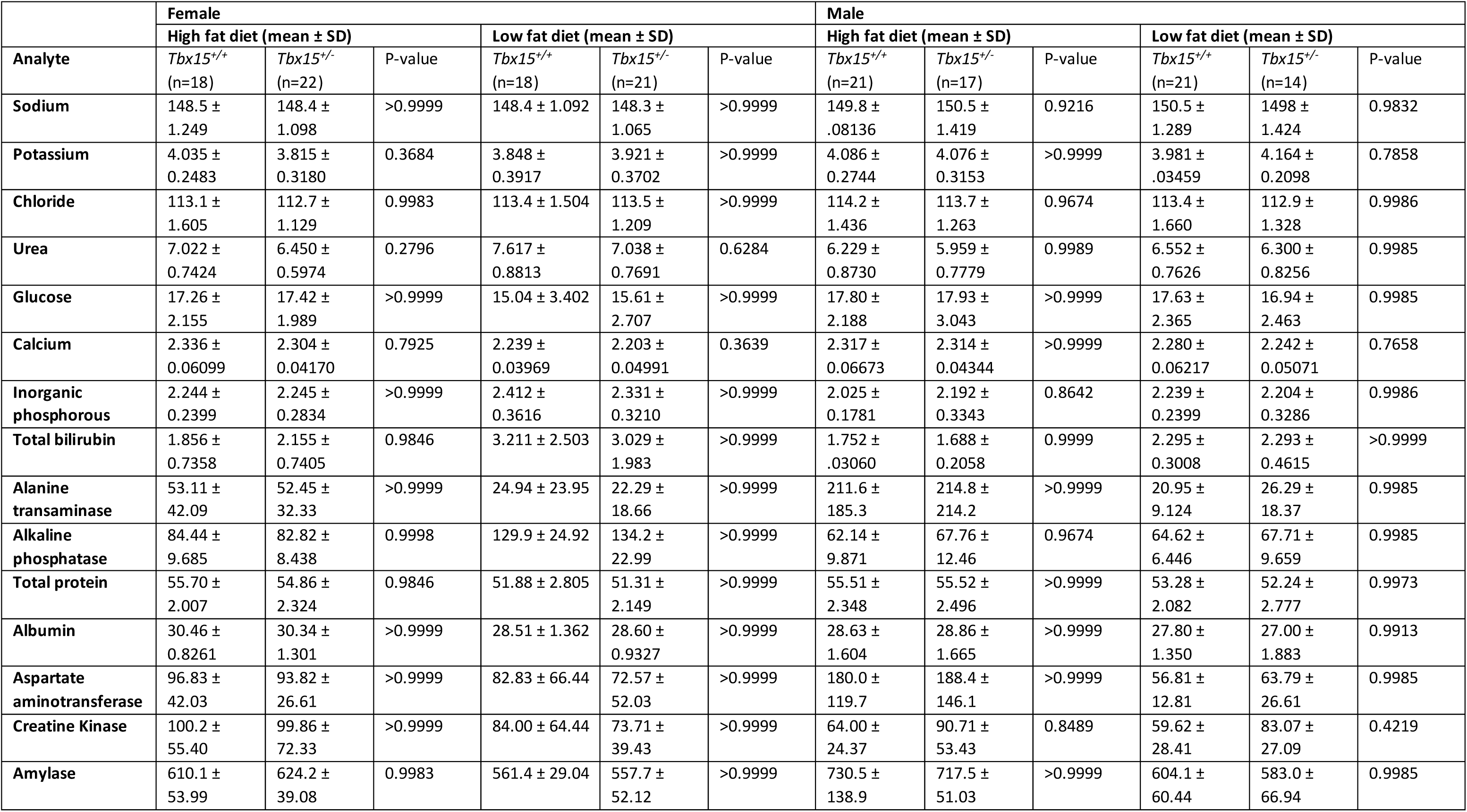
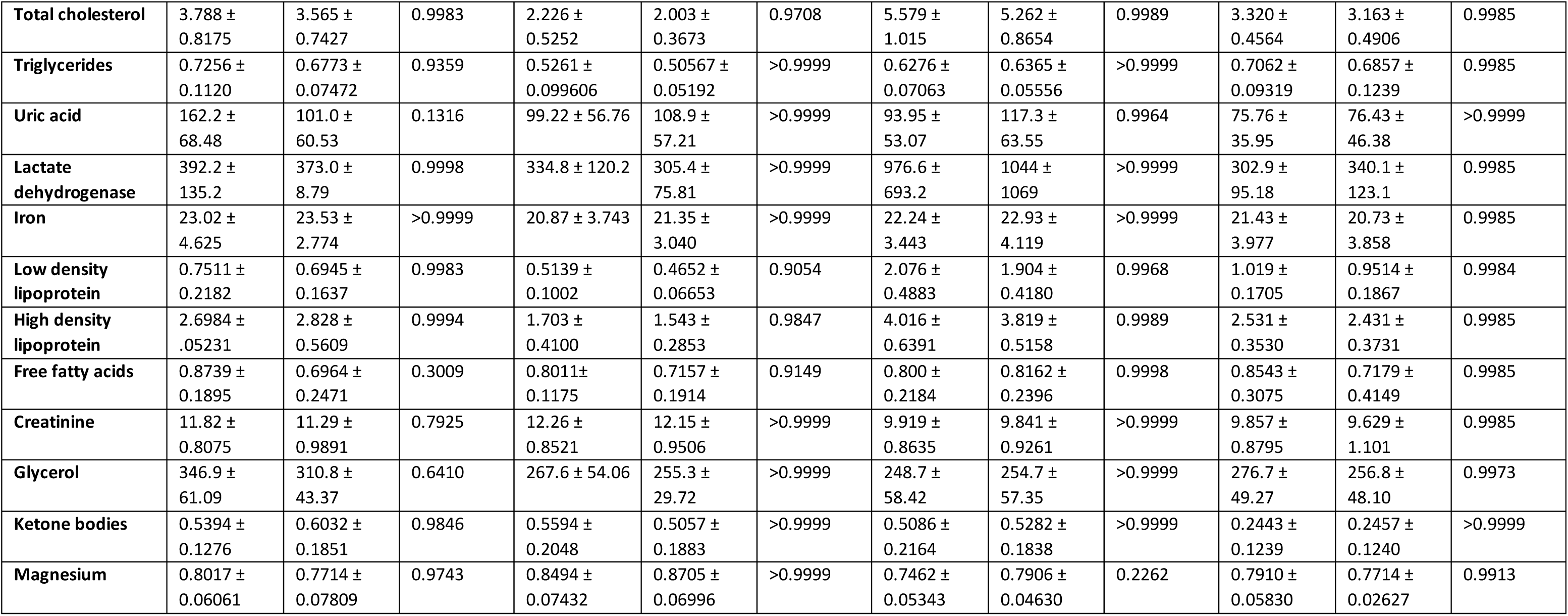
*Tbx15^+/-^*mice do not have altered plasma analyte levels at 24 weeks of age. Data are presented as mean ± standard deviation (n=14-22 per group). Statistical significance was calculated by repeated measures ANOVA for overall genotype and interaction (analyte x genotype) effects, with Holm-Šídák multiple comparisons test within each sex and diet. SD, standard deviation.

| Analyte | Female |  |  |  |  |  | Male |  |  |  |  |  |
| --- | --- | --- | --- | --- | --- | --- | --- | --- | --- | --- | --- | --- |
|  | High fat diet (mean ± SD) |  |  | Low fat diet (mean ± SD) |  |  | High fat diet (mean ± SD) |  |  | Low fat diet (mean ± SD) |  |  |
|  | *Tbx15*<sup>+/+</sup><br>(n=18) | *Tbx15*<sup>+/-</sup><br>(n=22) | P-value | *Tbx15*<sup>+/+</sup><br>(n=18) | *Tbx15*<sup>+/-</sup><br>(n=21) | P-value | *Tbx15*<sup>+/+</sup><br>(n=21) | *Tbx15*<sup>+/-</sup><br>(n=17) | P-value | *Tbx15*<sup>+/+</sup><br>(n=21) | *Tbx15*<sup>+/-</sup><br>(n=14) | P-value |
| **Sodium** | 148.5 ±<br>1.249 | 148.4 ±<br>1.098 | >0.9999 | 148.4 ± 1.092 | 148.3 ±<br>1.065 | >0.9999 | 149.8 ±<br>.08136 | 150.5 ±<br>1.419 | 0.9216 | 150.5 ±<br>1.289 | 1498 ±<br>1.424 | 0.9832 |
| **Potassium** | 4.035 ±<br>0.2483 | 3.815 ±<br>0.3180 | 0.3684 | 3.848 ±<br>0.3917 | 3.921 ±<br>0.3702 | >0.9999 | 4.086 ±<br>0.2744 | 4.076 ±<br>0.3153 | >0.9999 | 3.981 ±<br>.03459 | 4.164 ±<br>0.2098 | 0.7858 |
| **Chloride** | 113.1 ±<br>1.605 | 112.7 ±<br>1.129 | 0.9983 | 113.4 ± 1.504 | 113.5 ±<br>1.209 | >0.9999 | 114.2 ±<br>1.436 | 113.7 ±<br>1.263 | 0.9674 | 113.4 ±<br>1.660 | 112.9 ±<br>1.328 | 0.9986 |
| **Urea** | 7.022 ±<br>0.7424 | 6.450 ±<br>0.5974 | 0.2796 | 7.617 ±<br>0.8813 | 7.038 ±<br>0.7691 | 0.6284 | 6.229 ±<br>0.8730 | 5.959 ±<br>0.7779 | 0.9989 | 6.552 ±<br>0.7626 | 6.300 ±<br>0.8256 | 0.9985 |
| **Glucose** | 17.26 ±<br>2.155 | 17.42 ±<br>1.989 | >0.9999 | 15.04 ± 3.402 | 15.61 ±<br>2.707 | >0.9999 | 17.80 ±<br>2.188 | 17.93 ±<br>3.043 | >0.9999 | 17.63 ±<br>2.365 | 16.94 ±<br>2.463 | 0.9985 |
| **Calcium** | 2.336 ±<br>0.06099 | 2.304 ±<br>0.04170 | 0.7925 | 2.239 ±<br>0.03969 | 2.203 ±<br>0.04991 | 0.3639 | 2.317 ±<br>0.06673 | 2.314 ±<br>0.04344 | >0.9999 | 2.280 ±<br>0.06217 | 2.242 ±<br>0.05071 | 0.7658 |
| **Inorganic phosphorous** | 2.244 ±<br>0.2399 | 2.245 ±<br>0.2834 | >0.9999 | 2.412 ±<br>0.3616 | 2.331 ±<br>0.3210 | >0.9999 | 2.025 ±<br>0.1781 | 2.192 ±<br>0.3343 | 0.8642 | 2.239 ±<br>0.2399 | 2.204 ±<br>0.3286 | 0.9986 |
| **Total bilirubin** | 1.856 ±<br>0.7358 | 2.155 ±<br>0.7405 | 0.9846 | 3.211 ± 2.503 | 3.029 ±<br>1.983 | >0.9999 | 1.752 ±<br>.03060 | 1.688 ±<br>0.2058 | 0.9999 | 2.295 ±<br>0.3008 | 2.293 ±<br>0.4615 | >0.9999 |
| **Alanine transaminase** | 53.11 ±<br>42.09 | 52.45 ±<br>32.33 | >0.9999 | 24.94 ± 23.95 | 22.29 ±<br>18.66 | >0.9999 | 211.6 ±<br>185.3 | 214.8 ±<br>214.2 | >0.9999 | 20.95 ±<br>9.124 | 26.29 ±<br>18.37 | 0.9985 |
| **Alkaline phosphatase** | 84.44 ±<br>9.685 | 82.82 ±<br>8.438 | 0.9998 | 129.9 ± 24.92 | 134.2 ±<br>22.99 | >0.9999 | 62.14 ±<br>9.871 | 67.76 ±<br>12.46 | 0.9674 | 64.62 ±<br>6.446 | 67.71 ±<br>9.659 | 0.9985 |
| **Total protein** | 55.70 ±<br>2.007 | 54.86 ±<br>2.324 | 0.9846 | 51.88 ± 2.805 | 51.31 ±<br>2.149 | >0.9999 | 55.51 ±<br>2.348 | 55.52 ±<br>2.496 | >0.9999 | 53.28 ±<br>2.082 | 52.24 ±<br>2.777 | 0.9973 |
| **Albumin** | 30.46 ±<br>0.8261 | 30.34 ±<br>1.301 | >0.9999 | 28.51 ± 1.362 | 28.60 ±<br>0.9327 | >0.9999 | 28.63 ±<br>1.604 | 28.86 ±<br>1.665 | >0.9999 | 27.80 ±<br>1.350 | 27.00 ±<br>1.883 | 0.9913 |
| **Aspartate aminotransferase** | 96.83 ±<br>42.03 | 93.82 ±<br>26.61 | >0.9999 | 82.83 ± 66.44 | 72.57 ±<br>52.03 | >0.9999 | 180.0 ±<br>119.7 | 188.4 ±<br>146.1 | >0.9999 | 56.81 ±<br>12.81 | 63.79 ±<br>26.61 | 0.9985 |
| **Creatine Kinase** | 100.2 ±<br>55.40 | 99.86 ±<br>72.33 | >0.9999 | 84.00 ± 64.44 | 73.71 ±<br>39.43 | >0.9999 | 64.00 ±<br>24.37 | 90.71 ±<br>53.43 | 0.8489 | 59.62 ±<br>28.41 | 83.07 ±<br>27.09 | 0.4219 |
| **Amylase** | 610.1 ±<br>53.99 | 624.2 ±<br>39.08 | 0.9983 | 561.4 ± 29.04 | 557.7 ±<br>52.12 | >0.9999 | 730.5 ±<br>138.9 | 717.5 ±<br>51.03 | >0.9999 | 604.1 ±<br>60.44 | 583.0 ±<br>66.94 | 0.9985 |
| **Zolkiewski et al TBX15 transcription network in adipose** |  |  |  |  |  |  |  |  |  |  |  |  |
| **Total cholesterol** | 3.788 ±<br>0.8175 | 3.565 ±<br>0.7427 | 0.9983 | 2.226 ±<br>0.5252 | 2.003 ±<br>0.3673 | 0.9708 | 5.579 ±<br>1.015 | 5.262 ±<br>0.8654 | 0.9989 | 3.320 ±<br>0.4564 | 3.163 ±<br>0.4906 | 0.9985 |
| **Triglycerides** | 0.7256 ±<br>0.1120 | 0.6773 ±<br>0.07472 | 0.9359 | 0.5261 ±<br>0.099606 | 0.50567 ±<br>0.05192 | >0.9999 | 0.6276 ±<br>0.07063 | 0.6365 ±<br>0.05556 | >0.9999 | 0.7062 ±<br>0.09319 | 0.6857 ±<br>0.1239 | 0.9985 |
| **Uric acid** | 162.2 ±<br>68.48 | 101.0 ±<br>60.53 | 0.1316 | 99.22 ± 56.76 | 108.9 ±<br>57.21 | >0.9999 | 93.95 ±<br>53.07 | 117.3 ±<br>63.55 | 0.9964 | 75.76 ±<br>35.95 | 76.43 ±<br>46.38 | >0.9999 |
| **Lactate dehydrogenase** | 392.2 ±<br>135.2 | 373.0 ±<br>8.79 | 0.9998 | 334.8 ± 120.2 | 305.4 ±<br>75.81 | >0.9999 | 976.6 ±<br>693.2 | 1044 ±<br>1069 | >0.9999 | 302.9 ±<br>95.18 | 340.1 ±<br>123.1 | 0.9985 |
| **Iron** | 23.02 ±<br>4.625 | 23.53 ±<br>2.774 | >0.9999 | 20.87 ± 3.743 | 21.35 ±<br>3.040 | >0.9999 | 22.24 ±<br>3.443 | 22.93 ±<br>4.119 | >0.9999 | 21.43 ±<br>3.977 | 20.73 ±<br>3.858 | 0.9985 |
| **Low density lipoprotein** | 0.7511 ±<br>0.2182 | 0.6945 ±<br>0.1637 | 0.9983 | 0.5139 ±<br>0.1002 | 0.4652 ±<br>0.06653 | 0.9054 | 2.076 ±<br>0.4883 | 1.904 ±<br>0.4180 | 0.9968 | 1.019 ±<br>0.1705 | 0.9514 ±<br>0.1867 | 0.9984 |
| **High density lipoprotein** | 2.6984 ±<br>.05231 | 2.828 ±<br>0.5609 | 0.9994 | 1.703 ±<br>0.4100 | 1.543 ±<br>0.2853 | 0.9847 | 4.016 ±<br>0.6391 | 3.819 ±<br>0.5158 | 0.9989 | 2.531 ±<br>0.3530 | 2.431 ±<br>0.3731 | 0.9985 |
| **Free fatty acids** | 0.8739 ±<br>0.1895 | 0.6964 ±<br>0.2471 | 0.3009 | 0.8011±<br>0.1175 | 0.7157 ±<br>0.1914 | 0.9149 | 0.800 ±<br>0.2184 | 0.8162 ±<br>0.2396 | 0.9998 | 0.8543 ±<br>0.3075 | 0.7179 ±<br>0.4149 | 0.9985 |
| **Creatinine** | 11.82 ±<br>0.8075 | 11.29 ±<br>0.9891 | 0.7925 | 12.26 ±<br>0.8521 | 12.15 ±<br>0.9506 | >0.9999 | 9.919 ±<br>0.8635 | 9.841 ±<br>0.9261 | >0.9999 | 9.857 ±<br>0.8795 | 9.629 ±<br>1.101 | 0.9985 |
| **Glycerol** | 346.9 ±<br>61.09 | 310.8 ±<br>43.37 | 0.6410 | 267.6 ± 54.06 | 255.3 ±<br>29.72 | >0.9999 | 248.7 ±<br>58.42 | 254.7 ±<br>57.35 | >0.9999 | 276.7 ±<br>49.27 | 256.8 ±<br>48.10 | 0.9973 |
| **Ketone bodies** | 0.5394 ±<br>0.1276 | 0.6032 ±<br>0.1851 | 0.9846 | 0.5594 ±<br>0.2048 | 0.5057 ±<br>0.1883 | >0.9999 | 0.5086 ±<br>0.2164 | 0.5282 ±<br>0.1838 | >0.9999 | 0.2443 ±<br>0.1239 | 0.2457 ±<br>0.1240 | >0.9999 |
| **Magnesium** | 0.8017 ±<br>0.06061 | 0.7714 ±<br>0.07809 | 0.9743 | 0.8494 ±<br>0.07432 | 0.8705 ±<br>0.06996 | >0.9999 | 0.7462 ±<br>0.05343 | 0.7906 ±<br>0.04630 | 0.2262 | 0.7910 ±<br>0.05830 | 0.7714 ±<br>0.02627 | 0.9913 |

### RNA-sequencing of *Tbx15^-/-^*, *Tbx15^+/-^* and *Tbx15^+/+^* mouse adipose tissues

In order to identify the TBX15 transcriptional network in adipose cells, we next used RNA-sequencing of whole adipose tissues. RNA from iWAT was isolated from male and female 12-week-old mice of all three genotypes, maintained on standard chow diet. We found that in females a total of 3225 and 104 genes were significantly differentially expressed (FDR <0.05) in iWAT and BAT, respectively, between *Tbx15^-/-^* null and *Tbx15^+/+^* wildtype mice. In iWAT and BAT only 3 and 63 genes, respectively, were differentially expressed between *Tbx15^+/-^*heterozygous and *Tbx15^+/+^* mice. In iWAT, 897 genes were upregulated and 2328 genes were downregulated in female *Tbx15^-/-^* compared to *Tbx15^+/+^* mice. In male mice, we saw a similar pattern, albeit to a lesser degree, with 6 upregulated and 592 downregulated genes in iWAT of *Tbx15^-/-^* compared to *Tbx15^+/+^* mice (Figure 6A). Given the higher number of DEGs in iWAT we continued with functional analysis only in this tissue. For both female and male *Tbx15^-/-^* iWAT, there was an enrichment of DEGs in 55 and 24 pathways respectively, among the most significant associations were regulation of immune cell differentiation, type 1 diabetes mellitus, cytokine receptor interactions and chemokine signalling (q < 0.05; Figure 6B, C). Whilst several of these pathways were enriched in both sexes, this was observed to a lesser degree in males.

**Figure 6.**
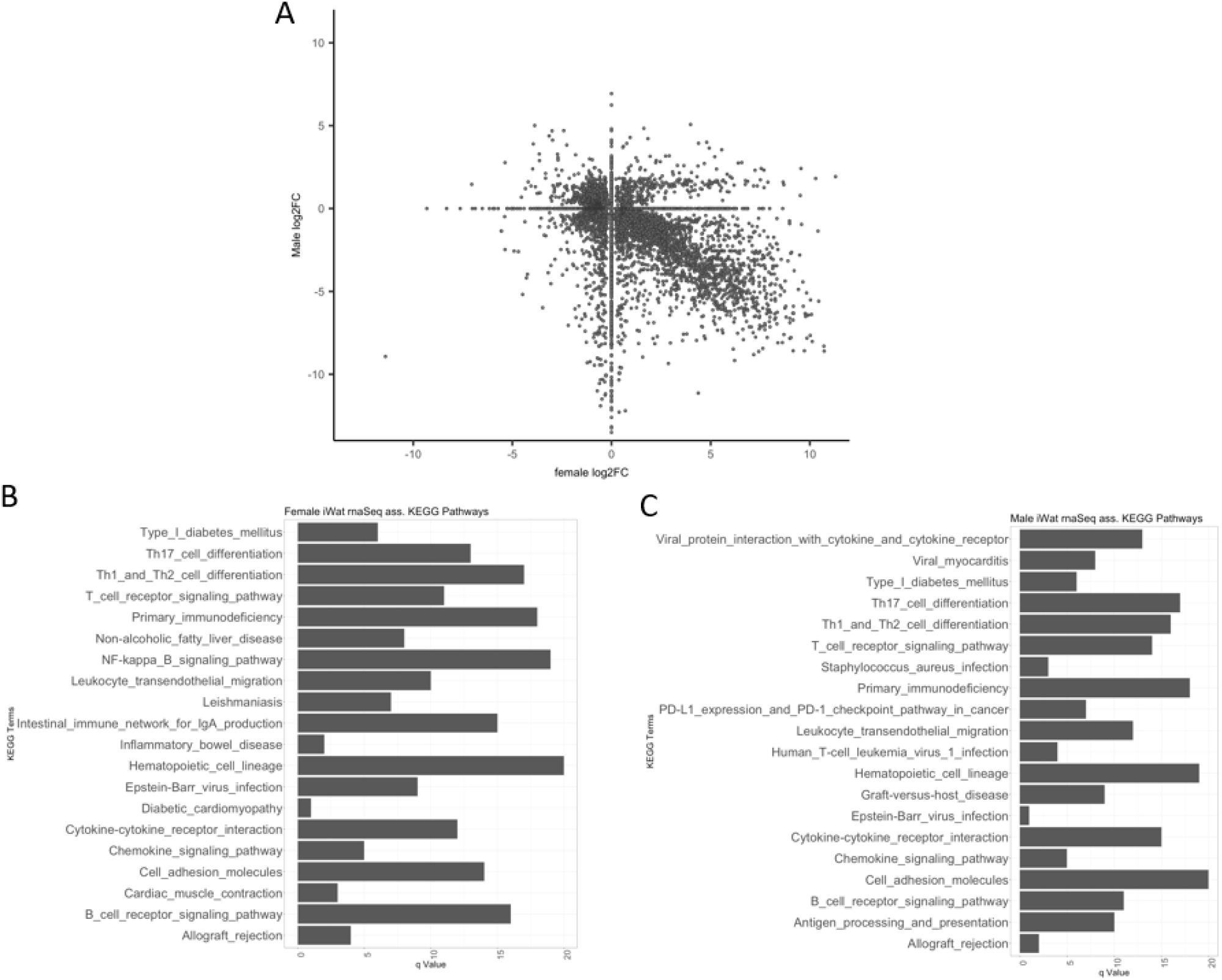
Sex differences in differentially expressed genes in iWAT of male and female *Tbx15^-/-^* mice. A. Scatterplot of log_2_ fold change (log2FC) gene expression differences in RNA isolated from iWAT of male and female *Tbx15^-/-^* vs *Tbx15^+/+^* controls. B-C. Over-represented KEGG pathways associated with differentially expressed genes (q<0.05) in *Tbx15^-/-^*female (B) and male (C) mice.

### ChIP-sequencing of 3T3-L1 mouse preadipocytes overexpressing TBX15

Next, we aimed to identify the genomic regions bound by the TBX15 transcription factor using the mouse (3T3-L1) preadipocyte cell line. We overexpressed HA-tagged human TBX15 (602 amino acids) in 3T3-L1 cells for 48 hours until post-confluence (the stage at which adipogenic differentiation can be initiated). An empty pCMV-HA vector was transfected as a negative control and cells were fixed for ChIP-sequencing analysis. . The majority of peaks were detected within 1500 bp of the genomic transcription start sites (TSS), with approximately twice as many peaks detected in regulatory regions such as enhancers rather than promoter regions (TSS; Figure 7**Error****! Reference source not found.**A and B). Target genes in the ChIP-sequencing analysis were enriched for 22 pathways, among the most significant pathways were thermogenesis, non-alcoholic fatty liver disease, insulin signalling and pathways of neurodegeneration (q<0.05; Figure 7**Error****! Reference source not found.**D).

**Figure 7.**
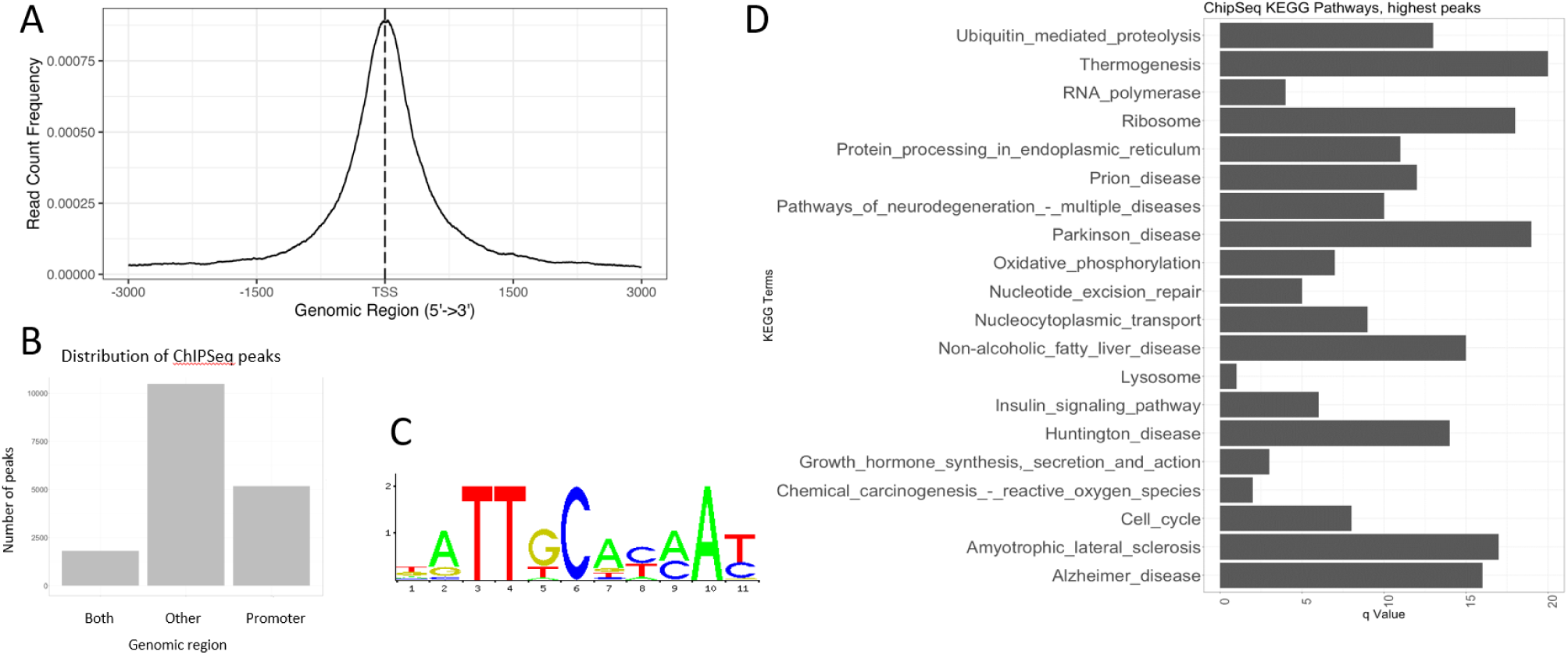
TBX15-HA acts primarily as an enhancer in 3T3-L1 preadipocyte cells. A. Distance (base pairs) of all annotated ChIP-sequencing TBX15 peaks from a transcription start site (TSS). B. Distribution of peaks in promoter and other regions of the genome. c) The T-box transcription factor binding motif was significantly over-represented at peaks (q<0.05). d). Over-represented KEGG pathways (q<0.05) associated with TBX15-HA ChIP-sequencing peaks.

### Generating a *critical set* of TBX15 transcriptional regulation in mouse reveals immune response and JAK/STAT signal pathway regulation

We then integrated the ‘omics datasets to define a core set of genes that are regulated by TBX15 and find pathways associated with these genes**Error! Reference source not found.**. We identified 52 common genes, which we termed the ‘*critical set*’ of TBX15 transcriptional regulation (Figure 8A, Table 3). The critical set are not significantly enriched in any functional pathways, however the pathways with the highest occurrence of genes compared to the reference genome are enriched in B- and T-cell receptor signalling, JAK-STAT signalling and haematopoietic cell lineage pathways suggesting a potential regulatory role for TBX15 in these pathways in adipose tissue (Figure 8**Error****! Reference source not found.**B).

**Figure 8.**
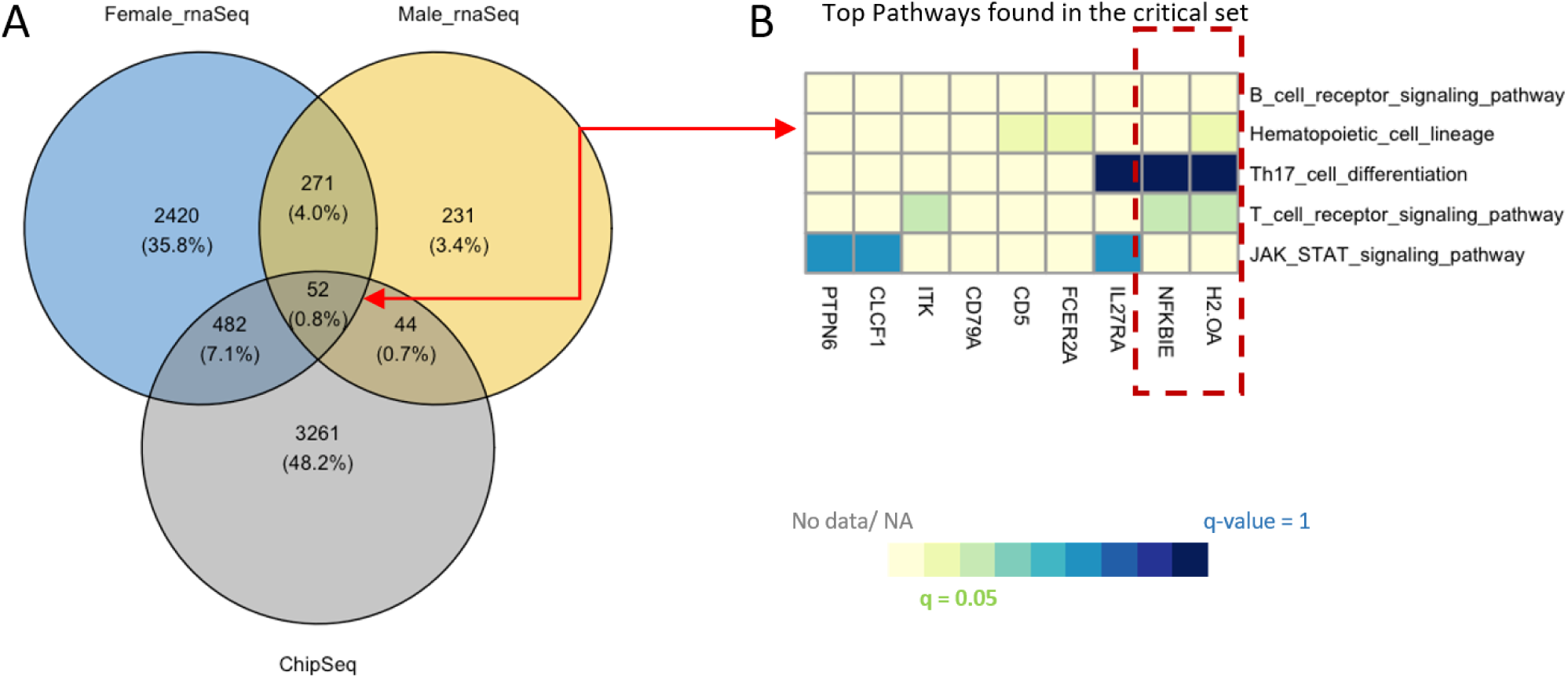
Integration of RNA-sequencing and ChIP-sequencing data reveals TBX15 regulates a critical set of 52 genes. A. Venn diagram of the number of differentially expressed genes (q<0.05) in female (blue circle) and male (yellow circle) *Tbx15^-/-^* iWAT versus *Tbx15^+/+^*controls and the number of peaks (q<0.05) bound by TBX15-HA (grey circle). 52 genes intersect all datasets called the “critical set”. B. Top 5 KEGG pathways associated with the genes found in the critical set. No KEGG pathways were significantly overrepresented in the critical set but two genes NFKBIE and H2-OA (red box) are also found in adipose associated pathways; Adipocytokine signalling pathway and Type I diabetes mellitus, respectively.

**Table 3.**
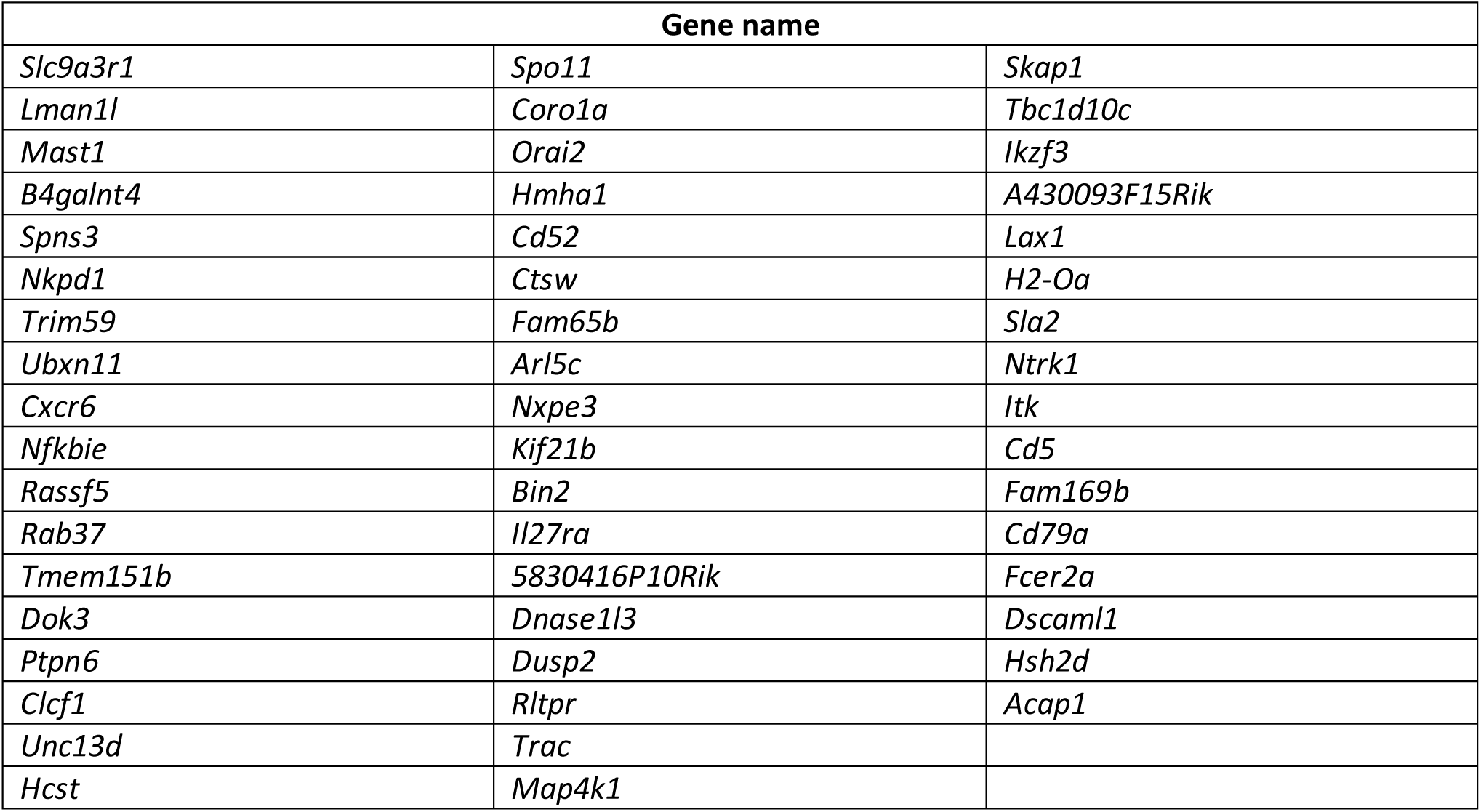
Integration of RNA-sequencing and ChIP-sequencing data reveals TBX15 regulates a critical set of 52 genes. List of 52 genes that intersect all datasets called the “critical set” from Figure 8A.

| Gene name |  |  |
| --- | --- | --- |
| *Slc9a3r1* | *Spo11* | *Skap1* |
| *Lman1l* | *Coro1a* | *Tbc1d10c* |
| *Mast1* | *Orai2* | *Ikzf3* |
| *B4galnt4* | *Hmha1* | *A430093F15Rik* |
| *Spns3* | *Cd52* | *Lax1* |
| *Nkpd1* | *Ctsw* | *H2-Oa* |
| *Trim59* | *Fam65b* | *Sla2* |
| *Ubxn11* | *Arl5c* | *Ntrk1* |
| *Cxcr6* | *Nxpe3* | *Itk* |
| *Nfkbie* | *Kif21b* | *Cd5* |
| *Rassf5* | *Bin2* | *Fam169b* |
| *Rab37* | *Il27ra* | *Cd79a* |
| *Tmem151b* | *5830416P10Rik* | *Fcer2a* |
| *Dok3* | *Dnase1l3* | *Dscaml1* |
| *Ptpn6* | *Dusp2* | *Hsh2d* |
| *Clcf1* | *Rltpr* | *Acap1* |
| *Unc13d* | *Trac* |  |
| *Hcst* | *Map4k1* |  |

## Discussion

In this work we demonstrate a role for TBX15 in adipose tissue biology and body composition. We found that global heterozygous *Tbx15* knockout was associated with reduced bodyweight in both male and female mice fed a LFD up to 25 weeks of age, coupled with lower fat mass in LFD female *Tbx15^+/-^* mice. These LFD *Tbx15^+/-^* mice had smaller WAT depots at 24-25 weeks of age. Interestingly, we did not detect a protective effect in *Tbx15^+/-^* mice from HFD-induced obesity, and female *Tbx15^+/-^* mice had marginally impaired glucose tolerance at 12 weeks. Differences in distribution of adipocyte cell area were detected in HFD , not LFD mice suggesting TBX15 may act through different mechanisms to regulate adipose tissue biology and body composition under different dietary challenges. Given these *in vivo* findings we assessed the TBX15 regulatory network using a combination of RNA-sequencing and ChIP-sequencing in iWAT and TBX15-over-expressing 3T3-L1 cell lines respectively. We identified thousands of DEGs in iWAT of female and male *Tbx15^-/-^* mice compared to *Tbx15^+/+^* littermates, with these genes enriched for immune response signalling pathways. When compared with TBX15 target genes identified by ChIP-sequencing we identified a “critical set” of 52 genes that we consider direct targets of TBX15 which were further enriched in immune cell receptor signalling pathways. Collectively our data indicate a role for TBX15 in adipose tissue biology, glucose tolerance and body composition. Further, our findings suggest a novel role for TBX15 regulating adipocyte-immune cell crosstalk in mature adipose tissues associated with obesity.

Previous *in vivo* studies identified an increased fat mass and glucose intolerance in heterozygous *Tbx15* knockout mice fed a standard RM3 diet ^18^ and adipose-specific *Tbx15* knockout mice fed a HFD ^19^ compared to their wild-type counterparts. Whilst we observed no whole body phenotypes or glucose tolerance difference in HFD *Tbx15^+/-^* mice in our study, we saw the converse in LFD *Tbx15^+/-^* mice which had less fat compared to *Tbx15^+/+^* littermates. These observed differences compared to previous work could be due to several factors including different regulatory effects of TBX15 under different dietary challenges, or in part due to the differences in metabolic phenotypes of the C57BL/6N and C57BL/6J backgrounds ^32^.

It is well established that *TBX15* is differentially expressed between subcutaneous and visceral adipose depots in both humans and mice^21,22,33,34^. Further, *Tbx15* has previously been implicated in the regulation of adipogenesis and browning of white adipose across a range of models ^19,21,22,35^. However, these data do not provide an unbiased view of the TBX15 regulatory network in adipose tissues and in some cases there is conflicting evidence for whether TBX15 acts as a transcriptional activator or repressor in adipogenesis. Here we present a comprehensive TBX15 regulatory network of over 3000 DEGs in iWAT, enriched for immune receptor signalling pathways. This, combined with identification of genomic TBX15 binding sites in 3T3-L1 preadipocyte cells highlights a core set of 52 genes bound and directly regulated by TBX15.

All 52 genes identified in the core set of direct TBX15 targets were downregulated in iWAT isolated from female *Tbx15^-/-^*mice compared to *Tbx15^+/+^*, suggesting in its natural state TBX15 acts as a transcriptional activator in adipose tissues. Actin cytoskeleton remodelling is a precursor for hypertrophic adipocyte expansion and is associated poor insulin tolerance *in vivo* ^36^. Downregulation of microtubule/cytoskeletal genes such as *Slc9a3r1, Mast1, Arl5c, Kif21b, Bin2, Rltpr* and *Tbc1d10c* could explain the differential adipocyte size distribution observed in *Tbx15^+/-^* mice. Indeed, *ARL5C* expression was positively correlated with accumulation of femoral fat mass in adipose-derived stem cells isolated from subcutaneous depots ^37^ and *BIN2* expression was increased in obese compared to lean individuals ^38^. The majority of the core 52 genes in our *critical set* are associated with leukocyte and lymphocyte activation, indicating a downregulation of the immune response in adipose tissues in *Tbx15^-/-^* mice. GWAS, epi-GWAS and exome sequencing have identified SNPs within several of these loci associated with a range of metabolic phenotypes for example *B4GALNT4* (T2DM, BMI and triglyceride levels), *TRIM59* (T2DM, body fat mass, body fat percentage and low-density lipoprotein [LDL] levels), *UBXN11* (fasting blood insulin levels and HOMA-IR) and *UNC13D* (Apolipoprotein A1 levels, BMI, HbA1c levels and cardiovascular disease) ^39–41^. *Acap1* directly binds to and promotes insulin-stimulated endocytic Glut4 recycling *in vitro* in 3T3-L1 preadipocytes ^42^. *LAX1,* predominantly involved in inhibition of MAPK signalling, is positively correlated with BMI and C-reactive protein expression in obese children ^43^. In NAFLD liver models, *TRIM59* overexpression promoted inflammation and steatosis, with *in vivo* knockdown of *Trim59* protecting against HFD-induced steatosis and expression of inflammatory cytokines TNF-α, IL-6 and IL-8 ^44^. In the context of TBX15, these data suggest that absence of *Tbx15* expression may reduce the cytoskeletal remodelling and recruitment of pro-inflammatory immune cells (M1 macrophages, Th1 cells) associated with hypertrophic adipose expansion that leads to obesity, resulting in the reduced bodyweights and fat mass phenotypes seen in *Tbx15^+/-^*mice.

This supports a potential role for TBX15 in regulating cytokine signalling in adipose tissues associated with expansion. It would appear that the downregulation of immune receptor signalling genes in iWAT of *Tbx15^-/-^*mice is reflected in the reduced fat mass seen in *Tbx15^+/-^* mice maintained on a LFD. Further research is required to elucidate the exact mechanisms through which TBX15 acts to regulate genes associated with altered adipose biology and body composition.

In conclusion, our data support a role for TBX15 in adipose tissue biology and body composition when faced with dietary challenge. Reduced *Tbx15* expression is associated with decreased fat mass and mildly impaired glucose tolerance in female mice. This could be attributed to the decreased immune receptor expression observed in iWAT with ablated *Tbx15* expression. This work highlights an important regulatory role for TBX15 in the adipocyte-immune cell crosstalk associated with obesity and type 2 diabetes mellitus and could provide new therapeutic targets in the future.

## Acknowledgements

The authors thank the staff of the Mary Lyon Centre and core services at MRC Harwell Institute for assistance with mouse studies. We thank the UKRI Medical Research Council (MC_U142661184 and MC_UP_2201/1) for supporting these studies. We thank the Oxford Genomics Centre at the Wellcome Centre for Human Genetics (Wellcome Trust 203141/Z/16/Z) for the generation and initial processing of the sequencing data. R. Dumbell is supported by Nottingham Trent University, the British Society for Neuroendocrinology, the Royal Society (RGS\R2\242302), and the Academy of Medical Sciences (AMS), the Wellcome Trust, the Government Department of Business, Energy and Industrial Strategy (BEIS), the British Heart Foundation and Diabetes UK, (SBF008\1169).

## Author contributions

**LZ**: Methodology, Investigation, Formal Analysis, Writing – Original Draft, Visualisation. **MS**: Methodology, Formal analysis, Visualisation. **JH:** Methodology. **LV:** Methodology. **EI:** Investigation. **LM:** Methodology, Investigation, Supervision. **MY:** Methodology, Supervision. **L. Beresford:** Investigation. **AR:** Investigation. **SH:** Investigation. **JH:** Investigation. **L. Bentley**: Methodology, Investigation, Supervision. **RDC**: Conceptualisation, Methodology, Writing – Review & Editing, Supervision, Funding Acquisition. **RD**: Conceptualisation, Methodology, Investigation, Writing – Review & Editing, Supervision.

